# Protein-Free ribosomal RNA scaffolds can assemble poly-lysine oligos from charged tRNA fragments

**DOI:** 10.1101/2021.01.12.426402

**Authors:** Doris Xu, Yuhong Wang

## Abstract

Ribosomal protein synthesis is a central process of the modern biological world. Because the ribosome contains proteins itself, it is very important to understand its precursor and evolution. Small ribozymes have demonstrated the principle of “RNA world” hypothesis, but protein free peptide ligase remains elusive. In this report, we have identified two fragments in the peptidyl transfer center that can synthesize a 9-mer poly-lysine in a solution contains Mg^2+^. This result is deduced from isotope-shifting in high resolution MS. To our best knowledge, this is the longest peptide oligo that can be synthesized by a pure ribozyme. Via single molecule FRET experiments, we have demonstrated the ligase mechanism was probably by substrate proximity via dimerization. We prospect that these RNA fragments can be useful to synthesize template free natural and non-natural peptides, to be model system for peptidyl transfer reaction mechanism and can shed light to the evolution of ribosome.

**Table of Content Graph:** 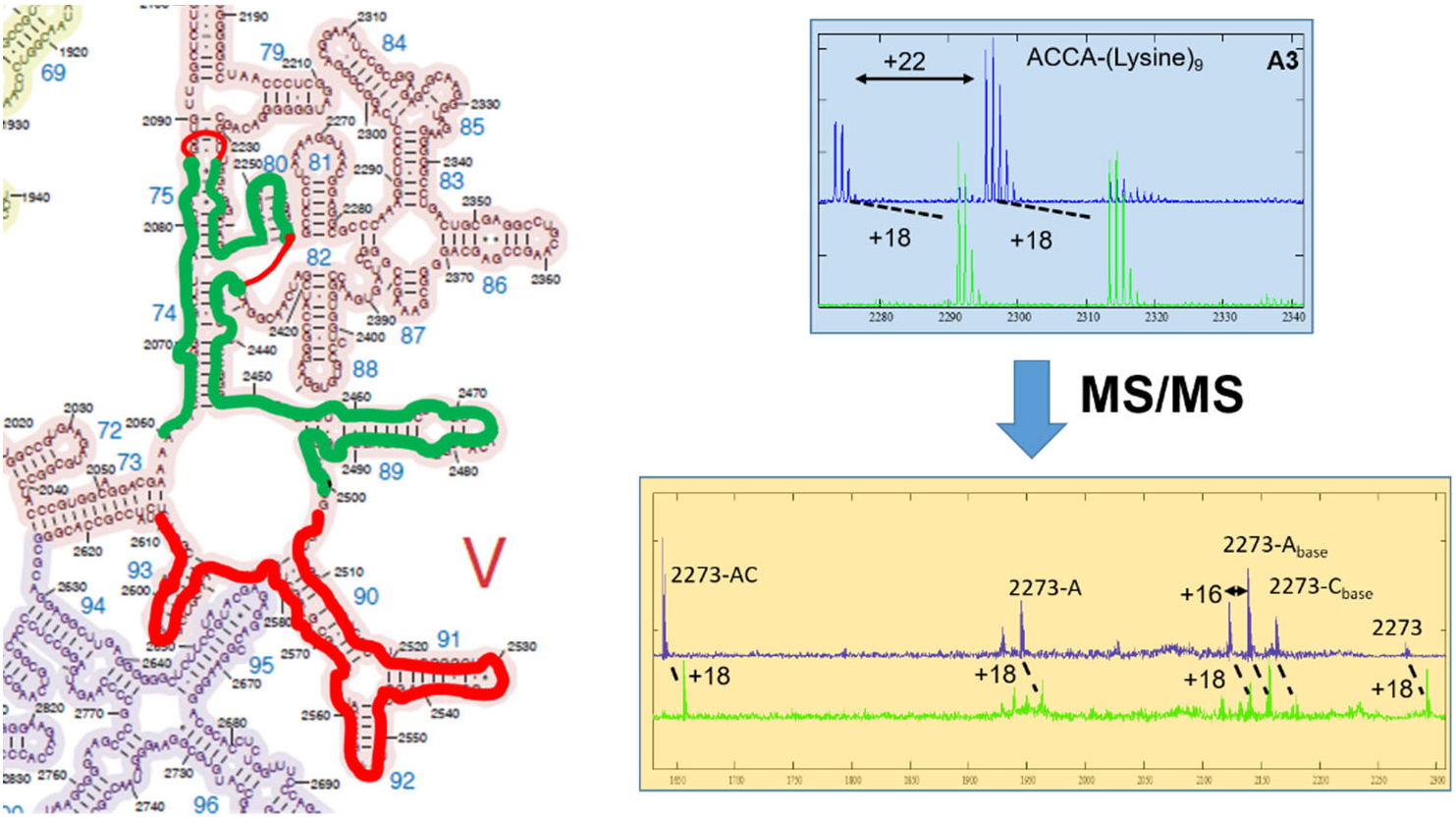

## Introduction

The ribosome is the universal molecular device that synthesizes all proteins in cells. These proteins are responsible for executing every major cellular function in the modern biological world. Ribosomes have two subunits (30S and 50S), each containing 20-50 proteins and 1-3 RNA polymers. Since modern ribosomes are made of the proteins that they synthesize, there must have existed protein-free ancestors that exhibited peptide ligase activity. The origin of peptide ligase is one of the most fundamental evolution questions to understand the transition from the “ancient RNA-” to the “modern protein-” worlds.[1,2] Before the divergence of the three life domains, the common ancestor of all organisms, LUCA (last universal common ancestor), contained nearly complete components for protein translation, implying that the development of the ribosome is an early event in the history of evolution. The lack of primitive intermediates also makes the origin of the ribosome elusive.[3]

Structural and biochemical assays have indicated that the peptidyl transfer center of the ribosome is the most ancient component and contains no protein.[4,5] However, the search for an RNA-only peptide ligase is still unsuccessful.[6,7] Nevertheless, very small non-rRNA ribozymes prepared through in vitro selection can catalyze peptide bond between short aminoacylated RNA oligos.[8,9] In addition, aminoacyl minihelix, which is more similar to current tRNA substrates, can form peptide bond with puromycin-containing oligos that complemented to the CCA sequence of the minihelix.[10] These results demonstrated the feasibility of RNA-only peptide ligase. However, truncated and in vitro selected constructs of longer RNA chains from the domain V of the large subunit rRNA 23S, in which the peptidyl transfer center resides,[11] were inactive of peptide bond formation. In the center of domain V, approximately 180 nucleotides formed a two-fold rotation symmetry where each half forms H-bond with the A- and P-site tRNAs, respectively.[12] These two fragments formed similar secondary and tertiary structures with little sequence homologs (Figure 1). Furthermore, residues G2553, G2252/G2253 (bacterial numbering) in the A-and P-loops formed H-bonds with the tRNAs to position them for optimized reaction orientation.[13,14]

**Figure 1.**
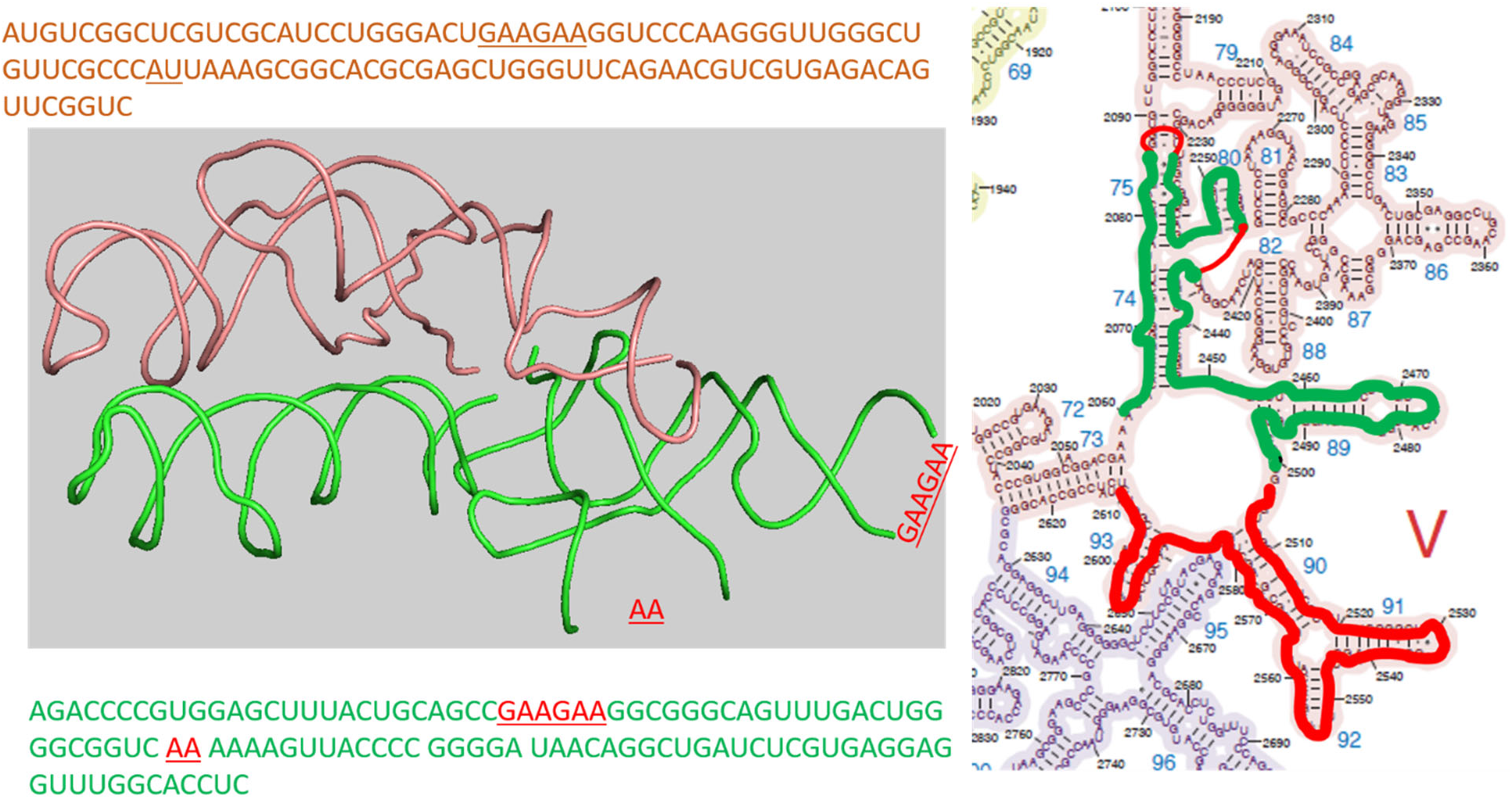
Short rRNA fragments at the peptidyl transfer with a 2-fold rotated symmetry that are outlined red and green. The left panel shows the 1^st^ and tertiary structures from 4wpo; the right panel shows the 2^nd^ structure.

Although proximity is not sufficient for peptide bond formation, it is a precondition.[10] Thus, we searched for the protein-free ligase using single molecule FRET detection. By attaching rRNA chains to a glass surface under a microscope, proximal binding of tRNAs is easy to detect without ambiguity. Surprisingly, we have found that a 108 nt rRNA scaffold (residues 2503-2610, named ptc1b and outlined red in Figure 1[15]) alone can bring two labeled tRNA 3’-fragements into very close distance, which is indicated by a FRET efficiency of 0.6. Further FRET experiments on labeled ptc1b suggested that closeness of the tRNA fragments was due to rRNA dimerization, which depended on Mg^2+^. By superimposing this rRNA piece into its symmetric counterpart in domain V (pdb ID 4wpo), we identified another rRNA scaffold (green outline in Figure 1, named ptc1a). It contains 2060 to 2501 fragments, with two stem loops truncated and filled with short RNA oligos guided by the superposition. Both sequences are shown in Figure 1. We then tested the ligase activities of the homo/hetero rRNA dimers with charged Lysine-tRNA^lysine^ after RNaseT digestion. Surprisingly, we identified formation of a 9-mer poly-lysine. This result is corroborated by mass-shifting of product incorporating N^15^,N^15^-labled lysine in high resolution Mass Spec. We found that rRNA alone can synthesize not just single peptide bonds but multiple ones, suggesting co-appearance and co-evolution of peptides and RNAs in the primordial world. The template-free protein synthesis seems to suggest a module-based evolution hypothesis, in which 3D structure interactions without sequence constrain drive the evolution.[16] The simple RNA scaffolds also provided an easy-to-manipulate model to study the chemical mechanism of the peptidyl transfer reactions, which remains a fundamental challenge.[17] Finally, we project that this simple template-free, proofreading-free protein polymerase will be useful to synthesize non-natural peptides for bioengineering applications.

## Materials and Methods

All chemicals were purchased from Millipore Sigma unless stated otherwise.

### RNA scaffolds

The 5’-biotinylated ptc1b RNA and ptc1a RNA molecules are purchased from IDTDNA. The 3’-ends are labeled with the “3’ EndTag™ DNA End Labeling System” from the Vector^@^Laboratories. The Cy3/Cy5-maleimide dyes are purchased from GE Life Sciences. The labeling efficiency is approximately 50% based on spectrophotometer absorptions.

### tRNA charging and Rnase T1 digestion

The tRNA^lysine^ powder is purchased from chemical-block.com and is charged with normal-(Millipore-Sigma), C^14^-(PerkinElmer), or N^15^,N^15^-(Millipore-Sigma) labeled lysine as described previously.[18] The charging efficiency is approximately 1000 pmol/A_260_ based on absorption and C^14^ radioactivity. Afterward, 4 A_260_ of charged tRNA^lysine^ (theoretical amount 1800 pmol/A_260_) is digested with 1,000 units of RNase T1 (thermofisher) under denature condition following the manufacture’s manual. The ACCCACCA-lysine piece is first phenol extracted and precipitated, and re-dissolved with water. Then it is enriched with the Monarch® RNA Cleanup kit. First, the RNA oligos > 25 nt is captured by the Monarch® binding column by adding 1x volume of ethanol. Then another 1x volume of ethanol is added to the run-through liquid, and loaded onto a new filter to bind the smaller pieces < 25 nt. After washing, the RNA-lysine piece is eluted with 50 μl of RNase-free water. The expected RNA-lysine pieces (ACCCACCA-lys^N14,N14^ and-lys^N15,N15^) are supported by high resolution MALDI MS (Mass Spectrometry facility, University of Alabama), and named “RNA-lys” (Figure 3A1).

### RNA-lysine labeling

The RNA-lysine piece was labeled in two positions: at the lysine moiety and the 5’-end of the RNA. <u>Labeling at the lysine moiety</u>. Cy3 or Cy5 NHS-dyes (GE Life Sciences) were dissolved in DMSO and added directly into the RNA-lysine solution mentioned above (final dye concentration ~ 500 μM from 10 mM stocks). The labeling reaction was incubated at 37°C for 2 hours. And the excess dye was removed via a G25 size-exclusive column. Then the Cy3-or Cy5-labeled RNA-lysine was precipitated and resuspended in 10-20 μl of RNase-free water. <u>Labeling at the 5’-end</u> was prepared with the “5’ EndTag™ DNA/RNA End Labeling System” from the Vector^@^Laboratories. The labeling efficiency of both position is approximately (0.6μM of dye)/(A_260_ of RNA).

### Single molecule FRET experiments

The total internal reflection fluorescence microscope (TIRF) was based on a Nikon Eclipse Ti2-E inverted microscope with two auto-turrets and two CMOS cameras as described before.[18] The top turret reflects the laser for TIRF illumination, and the bottom turret split the FRET signals into two CMOS cameras based on wavelengths. The sample holder was described previously.[19] For all measurements, sample concentrations are in the range of 10-100 nm. An oxygen scavenger cocktail (3 mg/mL glucose, 100 mg/mL glucose oxidase, 48 mg/mL catalase, and 2 mM trolox) was added to the channel before imaging to prevent photo bleaching.

## Results and Discussion

First, we have observed high FRET efficiency between Cy3/Cy5-labled RNA-lys in the presence of ptc1b. The reason ptc1b was chosen, instead of the total 180 nt from the literature,[12] was due to the complicated folding for longer RNAs. The sequence of ptc1b reproduced the same secondary structure as in the ribosome, while the 180 nt chain did not (Mfold Web Server). As shown in Figure 2A, the Cy3 and Cy5 dyes are located at the aminoacyl and the 5’-terminal of RAN-lys, respectively. 1 μM of ptc1b was incubated with 0.5 μM of Cy3/Cy5-labeld RNA-lys in the presence of 0 or 15 mM MgCl_2_. The mixture was incubated at 37°C for 10 min and diluted with either water or 15 mM MgCl_2_ solution at room temperature. This solution was loaded into the sample cell and attached to the surface via biotin-streptavidin interaction. Typical FRET images and representative time-lapse traces of donor and acceptor fluorescent intensities are shown in Figure S1. The FRET efficiencies are calculated as *I*_acceptor_/(*I*_donor_ + *I*_acceptor_), in which *I_xx_* is the fluorescence intensity in the donor or acceptor channel. In the presence of Mg^2+^, a FRET species centered at 0.62 was observed, which corresponds to 50 Å (R_0_ = 55 Å).[20] From the X-ray structure (4wpo), the amino acid of A-site tRNA to the 5’-of ACCCACCA at P-site (tRNA^lys^ sequence) is approximately 43 Å. This result suggested that the ptc1b can dimerize to mimic the two-fold symmetric peptidyl transfer center; and the tRNA fragments bind to the dimer in a similar way as tRNAs binding to the ribosome.

Secondly, the ptc1b/a dimerization hypothesis was tested. The ptc1b is labeled on the 3’-ends (C2610) with Cy3 and Cy5, respectively. Their dimerization is directly monitored with FRET. As predicted, FRET between Cy3-ptc1b and Cy5-ptc1b (Figure 2), exhibited a high FRET species centered at 0.60. Without Mg^2+^, this species disappeared (Figure S1), indicating that the dimerization also needs Mg^2+^. This FRET value corresponds to 51.4 Å, where the theoretical distance from the structure is about 23 Å (by aligning ptc1b to its hypothetical symmetric counterpart, insert of Figure 2A). This discrepancy indicates a looser structure in the RNA-only dimers than in the well-packed single-chain peptidyl transfer center. Variation of linker length, quantum yields and orientation may contribute to the difference as well. We then synthesized ptc1a molecule, in which most of the sequence were from rRNA, but two long stem loop structures were replaced with short oligos, based on its alignment with ptc1b (Figure S2).

**Figure 2.**
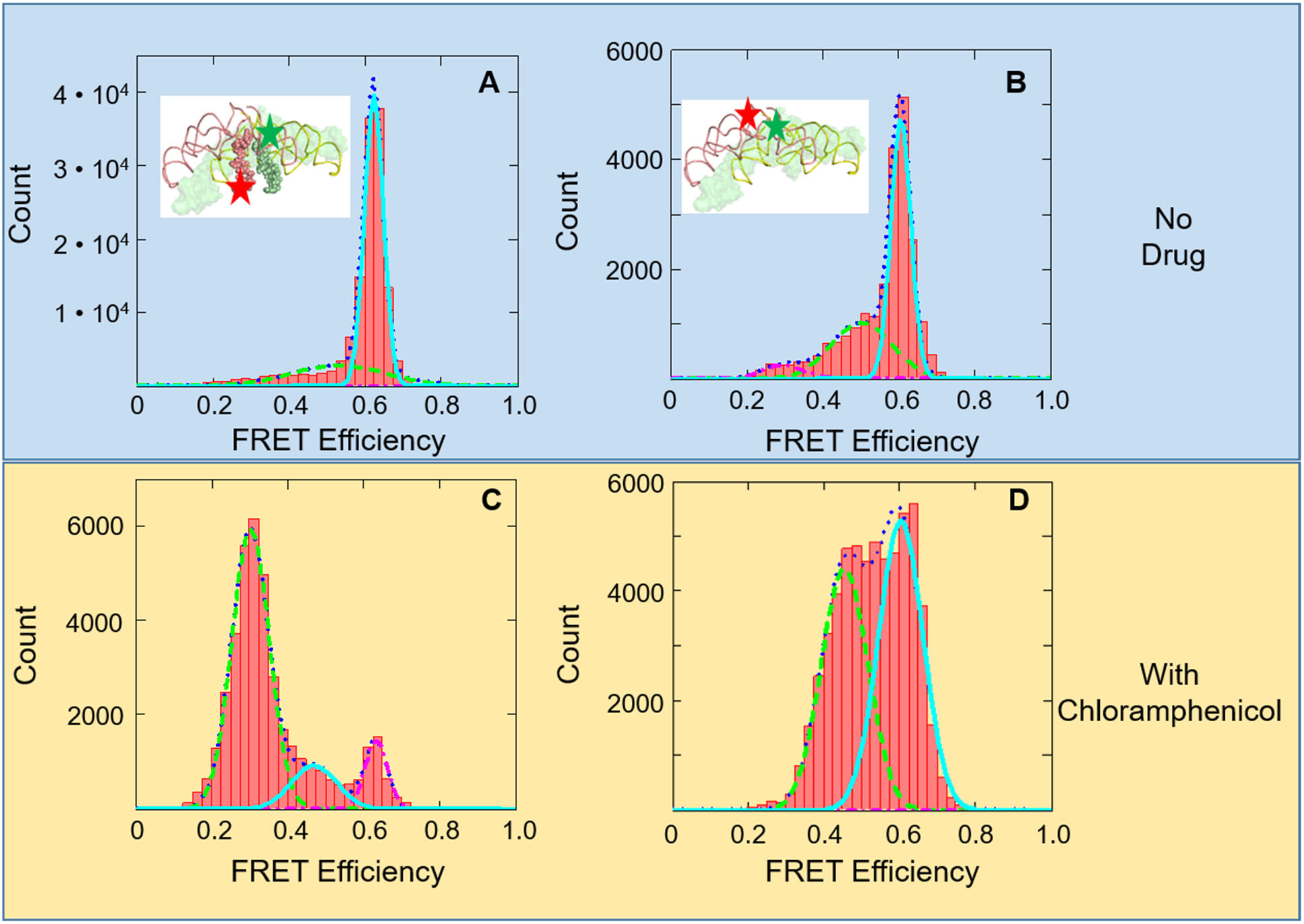
FRET efficiency histograms of tRNA (oligo)-tRNA (oligo) or rRNA-rRNA interactions. (A) FRET efficiency histogram of 5’- and aminoacyl labeled RNA-lys pieces. The relative Cy3/Cy5 positions are shown in the insert. (B) FRET efficiency histogram of 3’-labeled ptc1b. The relative Cy3/Cy5 positions are shown in the insert. (C) and (D) FRET efficiency histograms of (A) and (B) in the presence of 1 mM chloramphenicol, respectively.

The ptc1a piece was also labeled at the 3’-end, and hetero-dimer formation of ptc1a/b was detected. The FRET efficiencies are almost the same in the hetero dimer experiment (Figure S3). These high FRET species (ptc1b homo-dimer and the ptc1a/b hetero-dimer) are probably the reasons to bring the RNA-lys species together that generated the high FRET signals in Figure 2A.

Thirdly, the 0.62 FRET species between RNA-lys in Figue 2A was diminished by chloramphenicol, but the 0.60 FRET species between ptc1b homodimer in Figure 2B was not (Figure 2C and D). Chloramphenicol is an antibiotic that binds to the same location as the aminoacyl moiety of A-site tRNA.[21] If the prospected structures are correct in Figures 2A and 2B, then adding chloramphenicol should interfere the high FRET species in 2A, but not in 2B.

This is because chloramphenicol directly competes the binding pocket with RNA-lys, but does not compete with the ptc1a/b dimerization without the short tRNA fragments.

We observed this effect in Figures 2C and D. The 0.62 FRET species diminished while a 0.3 FRET species emerged in the RNA-lys experiments, indicating inhibition of RNA-lys binding. Meanwhile, the 0.6 FRET species remained unchanged in the ptc1a/b dimerization experiments. Furthermore, the small peak at 0.5 FRET species in Figure 2B is significantly larger when drug is present in Figure 2D, which means chloramphenicol may have induced more dimers in a more loose structure. Figures 2C and 2D suggest that the structural assignments of the FRET species in Figures 2A and 2B are reasonable.

Fourthly, we have found that a 9-mer poly-lysine oligo is formed by the ptc dimers. In these experiments, 1 μM of ptc1b homo-dimer or ptc1a/b hetero-dimer was incubated with 15 mM MgCl_2_ and 1 μM of RNA-lys, in which normal or N^15^,N^15^-labeled lysine was used. The mixtures were incubated at 37°C for 1hr or 3 hrs (no difference was detected in mass spectrometry analysis), run through a G25 desalting column, and analyzed with high resolution MALDI (all the exact masses are shown in Table S1). Multiple mass peaks exhibited +18 shift in comparison of the N^14^ (blue trace) to the N^15^ (green trace) labeled experiments, showing incorporation of 9 lysine residues (2 Da mass shift per residue). Tandem MS/MS spectra were obtained on the 2273 peak and its +18 shifting peak in the normal and N^15^, N^15^-labeled experiments, respective (Figure 3B). Again, these spectra showed consistent +18 shifting at multiple positions, indicating there were 9-mer lysine in the 2273/2291 peaks. Mass calculation indicated that these peaks correspond to CCA-K_9_ (Figure 3A2), ACCA-K_9_ (Figure 3A3), CCACCA-K_9_ (Figure 3A4), CCCACCA-K_9_ (Figure 3A5). For the MS/MS of 2273 peak, species losing A or C base on the riboses and losing complete A- and AC-nucleotides are observed. Although the starting material was confirmed to be ACCCACCA-K (Figure 3A1, material and methods), we did not observe the product of 9-mer poly-lysine on the 8-nt fragment. It is probably due to the loss of the terminal A residue, either during the reaction incubation or MALDI measurements.[22]

**Figure 3.**
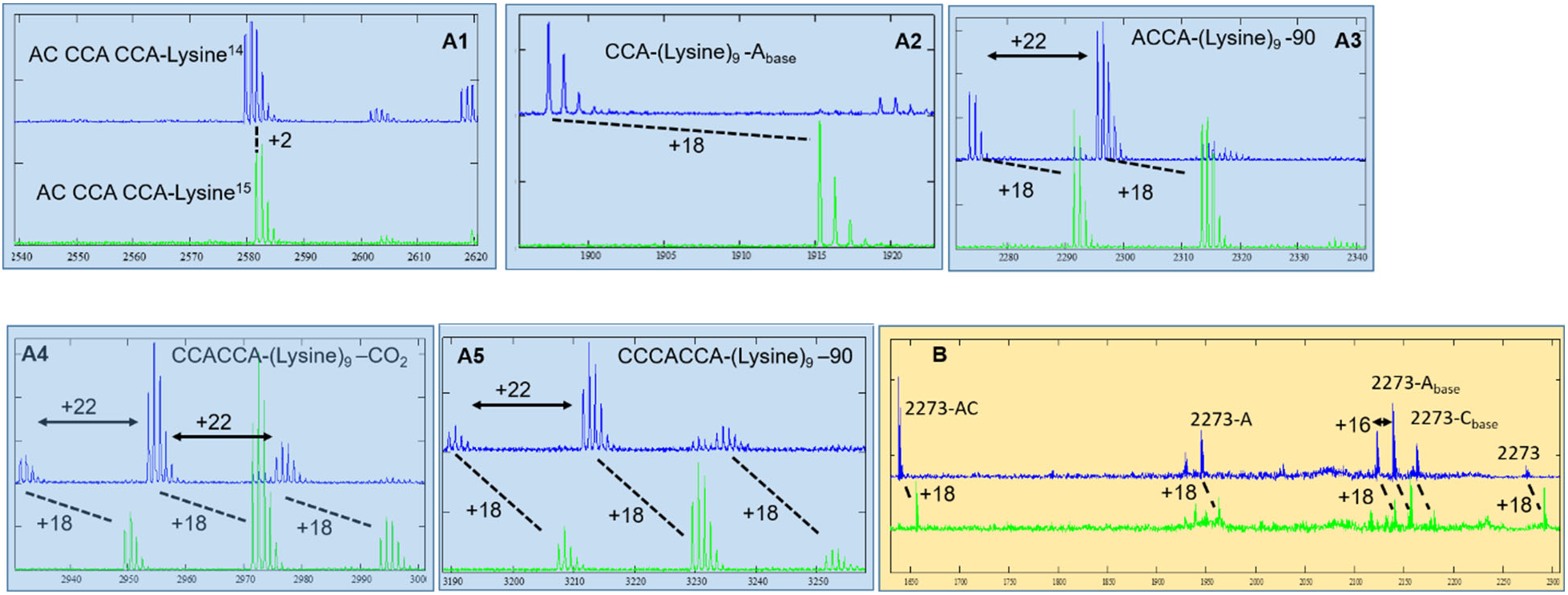
High resolution MS of poly-lysine of N^14^,N^14^- (blue trace) and N^15^,N^15^- (green trace) labeled lysine monomers. Shifting of 18 suggests 9 lysine residues. The exact masses and assignments are shown in Tables S1 and S2. (A1-5) Mass spectrum of lysine monomer and oligos. (B) Tandem MS/MS on 2273 and 2291 peaks.

In summary, short rRNA fragments, containing the same sequences as those at the peptidyl transfer center, dimerized and brought tRNA fragments to proximity. High resolution mass spectrometry analysis indicated that these dimers have synthesized not one but eight peptide bonds to form a 9-mer RNA-K_9_ molecule out of the Lysine-charged tRNA fragment (ACCCACCA-K). Poly-lysine has shown to adopt β-sheets structure, which intercalates well into the double helix grooves.[16,23] The ancient molecules tRNA synthetases and ribosome reserved these interactions. The tRNA synthetases depend on their β-sheets structures to charging tRNAs,[24] while some ribosomal proteins bind to rRNA’s grooves. For example L15 and S11 proteins demonstrated such interaction (Figure S4). This relatively long oligo synthesis we have observed suggests that primitive peptide ligase may have higher capacity than we thought. Therefore, a peptide/nucleic acid co-evolution route is possible, as suggested in the literature.[25] Theoretic study has suggested that optimizing the interaction between RNA’s double helix and peptide’s β-sheet could drive the evolution of both species without sequencing fidelity.[16] Indeed, diverse RNA sequences can form similar structural modules.[26,27] The ptc1a and 1b molecules in this report are examples. Alternatively, the discovery of RNA-only self-replicators strongly supports the “RNA world” theory, which is another possibility.[28,29]

One potential application of our finding is to mutate these simple rRNA fragments to test hypothetical catalytic residues at the peptidyl center. Although a major catalytic mechanism of the ribosome is to position the substrates at proximity, it remains unclear whether any chemical catalytic mechanism is involved. Several highly conserved residues have been tested, but none seem indispensable.[30-32] Using this short rRNA fragment that can be readily mutated and transcribed in vitro, we expect that some chemical mechanism of the peptidyl transfer reaction will be revealed, although this system is overly simplified and primitive.

We also propose that this simple peptide ligase can be a starting point in bioengineering to synthesize non-natural peptides without mRNA templates, which is difficult to incorporate by the native ribosome and is seriously pursued for drug development.[33] Synthesis of a 9-mer peptide oligo is a promising starting point to engineer a more powerful peptide ligase to make longer peptide chains without overcoming the rigorous selection and proofreading steps of the native ribosome. Therefore, unnatural/mixed peptide chains can be synthesized in the future. Acknowledgements

This research is supported by the the Welch foundation (grant number E-1721) and NIGMS (grant number 1 R01 GM111452) to Y.W.

We thank Dr. Qiaoli Liang of University of Alabama, Department of Chemistry for her excellent collection of the mass spectra.

## References

[1] F.H. Crick, The origin of the genetic code, J Mol Biol 38 (1968) 367–379.https://doi.org/10.1016/0022-2836(68)90392-6.

[2] W. Gilbert, Origin of life: The RNA world, Nature 319 (1986) 618–618. https://doi.org/10.1038/319618a0.

[3] G.E. Fox, Origin and evolution of the ribosome, Cold Spring Harb Perspect Biol 2 (2010) a003483. https://doi.org/10.1101/cshperspect.a003483.

[4] P. Nissen, J. Hansen, N. Ban, P.B. Moore, T.A. Steitz, The structural basis of ribosome activity in peptide bond synthesis, Science 289 (2000) 920–930. https://doi.org/10.1126/science.289.5481.920.

[5] C. Hsiao, T.K. Lenz, J.K. Peters, P.Y. Fang, D.M. Schneider, E.J. Anderson, T. Preeprem, J.C. Bowman, E.B. O’Neill, L. Lie, S.S. Athavale, J.J. Gossett, C. Trippe, J. Murray, A.S. Petrov, R.M. Wartell, S.C. Harvey, N.V. Hud, L.D. Williams, Molecular paleontology: a biochemical model of the ancestral ribosome, Nucleic Acids Res 41 (2013) 3373–3385. https://doi.org/10.1093/nar/gkt023.

[6] H.F. Noller, V. Hoffarth, L. Zimniak, Unusual resistance of peptidyl transferase to protein extraction procedures, Science 256 (1992) 1416–1419. https://doi.org/10.1126/science.1604315.

[7] P. Khaitovich, T. Tenson, A.S. Mankin, R. Green, Peptidyl transferase activity catalyzed by protein-free 23S ribosomal RNA remains elusive, RNA 5 (1999) 605–608. https://doi.org/10.1017/s1355838299990295.

[8] B. Zhang, T.R. Cech, Peptidyl-transferase ribozymes: trans reactions, structural characterization and ribosomal RNA-like features, Chem Biol 5 (1998) 539–553. https://doi.org/10.1016/s1074-5521(98)90113-2.

[9] B. Zhang, T.R. Cech, Peptide bond formation by in vitro selected ribozymes, Nature 390 (1997) 96–100. https://doi.org/10.1038/36375.

[10] K. Tamura, P. Schimmel, Oligonucleotide-directed peptide synthesis in a ribosome- and ribozyme-free system, Proc Natl Acad Sci U S A 98 (2001) 1393–1397.https://doi.org/10.1073/pnas.98.4.1393.

[11] R.M. Anderson, M. Kwon, S.A. Strobel, Toward ribosomal RNA catalytic activity in the absence of protein, J Mol Evol 64 (2007) 472–483.https://doi.org/10.1007/s00239-006-0211-y.

[12] I. Agmon, A. Bashan, R. Zarivach, A. Yonath, Symmetry at the active site of the ribosome: structural and functional implications, Biol Chem 386 (2005) 833–844.https://doi.org/10.1515/BC.2005.098.

[13] D.F. Kim, R. Green, Base-pairing between 23S rRNA and tRNA in the ribosomal A site, Mol Cell 4 (1999) 859–864.https://doi.org/10.1016/s1097-2765(00)80395-0.

[14] R.R. Samaha, R. Green, H.F. Noller, A base pair between tRNA and 23S rRNA in the peptidyl transferase centre of the ribosome, Nature 377 (1995) 309–314.https://doi.org/10.1038/377309a0.

[15] H.F. Noller, J. Kop, V. Wheaton, J. Brosius, R.R. Gutell, A.M. Kopylov, F. Dohme, W. Herr, D.A. Stahl, R. Gupta, C.R. Waese, Secondary structure model for 23S ribosomal RNA, Nucleic Acids Res 9 (1981) 6167–6189.https://doi.org/10.1093/nar/9.22.6167.

[16] C.W. Carter, Jr., J. Kraut, A proposed model for interaction of polypeptides with RNA, Proc Natl Acad Sci U S A 71 (1974) 283–287.https://doi.org/10.1073/pnas.71.2.283.

[17] R. Green, J.R. Lorsch, The path to perdition is paved with protons, Cell 110 (2002) 665–668.https://doi.org/10.1016/s0092-8674(02)00965-0.

[18] M.E. Altuntop, C.T. Ly, Y. Wang, Single-molecule study of ribosome hierarchic dynamics at the peptidyl transferase center, Biophys J 99 (2010) 3002–3009.https://doi.org/10.1016/j.bpj.2010.08.037.

[19] R. Lin, Y. Wang, Automated smFRET microscope for the quantification of label-free DNA oligos, Biomed Opt Express 10 (2019) 682–693.https://doi.org/10.1364/BOE.10.000682.

[20] T. Ha, I. Rasnik, W. Cheng, H.P. Babcock, G.H. Gauss, T.M. Lohman, S. Chu, Initiation and re-initiation of DNA unwinding by the Escherichia coli Rep helicase, Nature 419 (2002) 638–641.https://doi.org/10.1038/nature01083.

[21] D. Bulkley, C.A. Innis, G. Blaha, T.A. Steitz, Revisiting the structures of several antibiotics bound to the bacterial ribosome, Proc Natl Acad Sci U S A 107 (2010) 17158–17163.https://doi.org/10.1073/pnas.1008685107.

[22] J.C. Joyner, K.D. Keuper, J.A. Cowan, Analysis of RNA cleavage by MALDI-TOF mass spectrometry, Nucleic Acids Res 41 (2013) e2.https://doi.org/10.1093/nar/gks811.

[23] W. Dzwolak, R. Ravindra, C. Nicolini, R. Jansen, R. Winter, The diastereomeric assembly of polylysine is the low-volume pathway for preferential formation of beta-sheet aggregates, J Am Chem Soc 126 (2004) 3762–3768.https://doi.org/10.1021/ja039138i.

[24] L. Ribas de Pouplana, P. Schimmel, Two classes of tRNA synthetases suggested by sterically compatible dockings on tRNA acceptor stem, Cell 104 (2001) 191–193.https://doi.org/10.1016/s0092-8674(01)00204-5.

[25] B. Piette, J.G. Heddle, A Peptide-Nucleic Acid Replicator Origin for Life, Trends Ecol Evol 35 (2020) 397–406.https://doi.org/10.1016/j.tree.2020.01.001.

[26] Z. Miao, E. Westhof, RNA Structure: Advances and Assessment of 3D Structure Prediction, Annu Rev Biophys 46 (2017) 483–503.https://doi.org/10.1146/annurev-biophys-070816-034125.

[27] T. Hermann, E. Westhof, Non-Watson-Crick base pairs in RNA-protein recognition, Chem Biol 6 (1999) R335–343.https://doi.org/10.1016/s1074-5521(00)80003-4.

[28] D.S. Wilson, J.W. Szostak, In vitro selection of functional nucleic acids, Annu Rev Biochem 68 (1999) 611–647.https://doi.org/10.1146/annurev.biochem.68.1.611.

[29] K.F. Tjhung, M.N. Shokhirev, D.P. Horning, G.F. Joyce, An RNA polymerase ribozyme that synthesizes its own ancestor, Proc Natl Acad Sci U S A 117 (2020) 2906–2913.https://doi.org/10.1073/pnas.1914282117.

[30] G.W. Muth, L. Chen, A.B. Kosek, S.A. Strobel, pH-dependent conformational flexibility within the ribosomal peptidyl transferase center, RNA 7 (2001) 1403–1415.https://doi.org/.

[31] E.M. Youngman, J.L. Brunelle, A.B. Kochaniak, R. Green, The active site of the ribosome is composed of two layers of conserved nucleotides with distinct roles in peptide bond formation and peptide release, Cell 117 (2004) 589–599.https://doi.org/10.1016/s0092-8674(04)00411-8.

[32] N. Polacek, M. Gaynor, A. Yassin, A.S. Mankin, Ribosomal peptidyl transferase can withstand mutations at the putative catalytic nucleotide, Nature 411 (2001) 498–501.https://doi.org/10.1038/35078113.

[33] D.D. Young, P.G. Schultz, Playing with the Molecules of Life, ACS Chem Biol 13 (2018) 854–870.https://doi.org/10.1021/acschembio.7b00974.

