## supplemental file for "Protein-Free ribosomal RNA scaffolds can assemble poly-lysine oligos from charged tRNA fragments"

Table of contents:

| Content | page |
|---|---|
| Title | 1 |
| Figure S1 | 2 |
| Figure S2 | 3 |
| Figure S3 | 4 |
| Figure S4 | 5 |
| Table S1 | 6 |
| Table S2 | 6 |
| Reference | 7 |

Figure S1

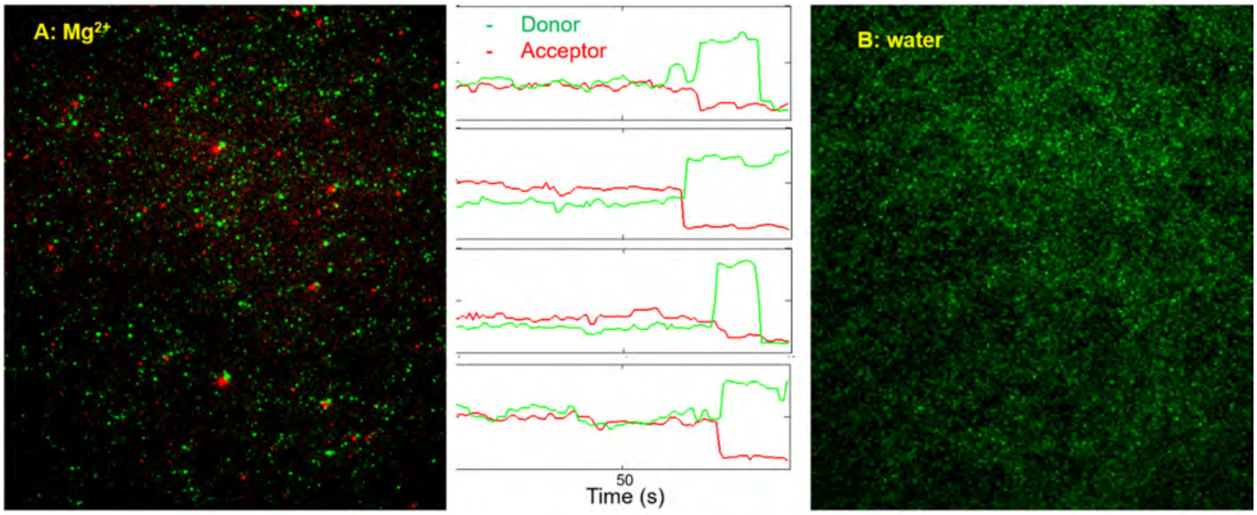


**Figure S1.** Examples of smFRET images and fitted fluorescence intensity time-lapse traces: (A) in the presence (high FRET formed) and (B) absence of $Mg^{2+}$ (no FRET). The Cy3/Cy5 labels are shown in Figure 2. There are two type of Cy3/Cy5 experiments, both showed the similar images as above.

Figure S2.

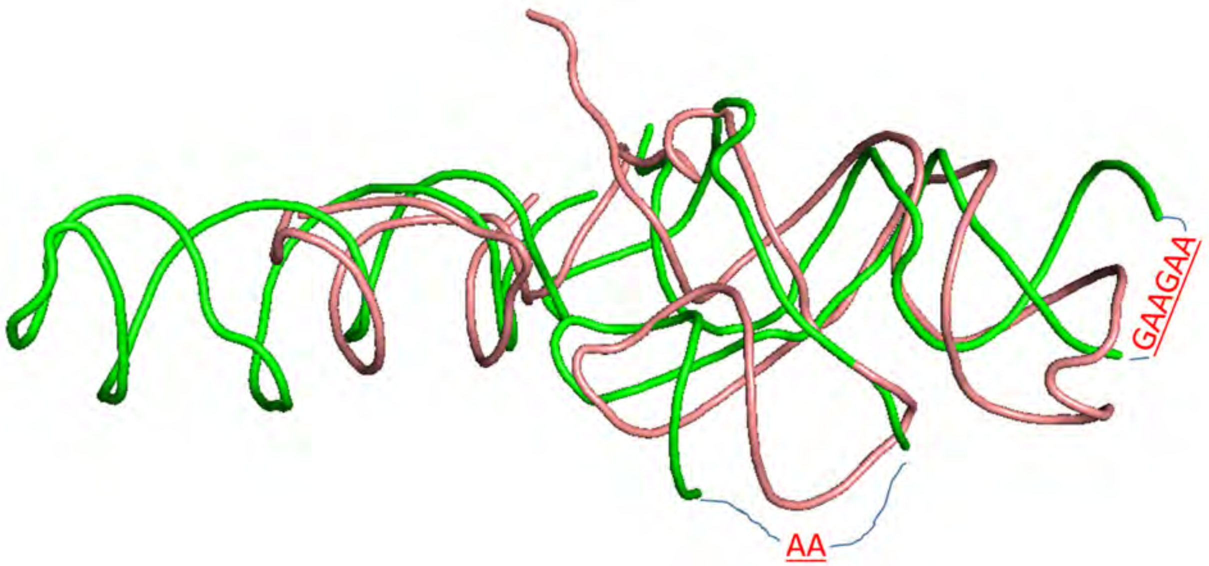


**Figure S2.** Alignment of ptc1b (red) onto ptc1a (green) fragments. Two long stems in ptc1a are truncated and replaced with the sequence of ptc1b.

Figure S3

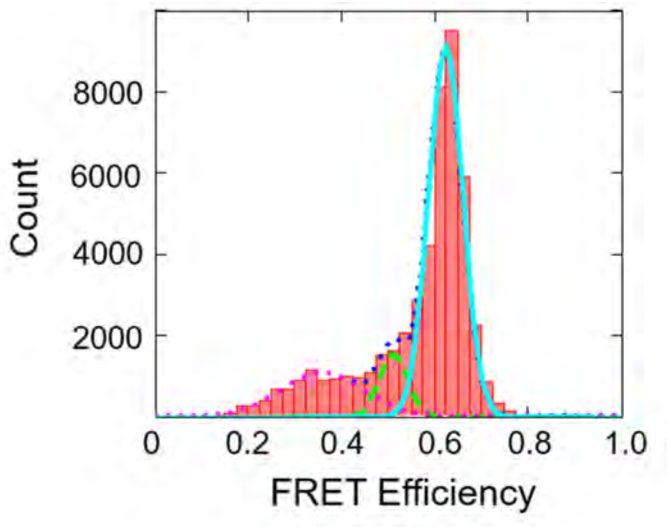


**Figure S3.** FRET efficiency histogram of heterodimer between ptc1a-Cy3 and ptc1b-Cy5. Both fragments are labeled at 3'-end. But one is labeled with Cy3 and the other one is labeled with Cy5.

Figure S4

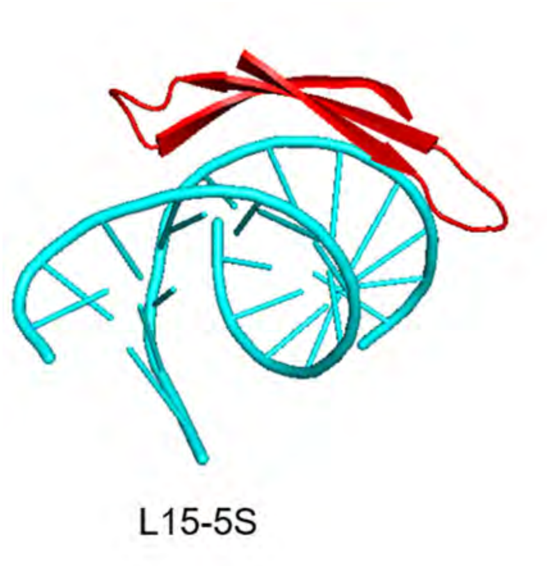


**Figure S4.** RNA duplex and protein b-sheet interactions between L15 and 5S; and between S11 and 16S. These 3D interactions are similar regardless of little sequence resemblance.

Table S1. Theoretical and measured Mass and analysis. Except for the substrates, only the $N^{14}$, $N^{14}$-lysine products are shown. The $N^{15}$,$N^{15}$-lysine products shift +18 in all the mass spectrum.

| species | Theoretical Mass (M+1) | Measured Mass | Difference | Assignment of difference (theoretical Mass) |
|---|---|---|---|---|
| ACCCACCA-$K^{N14,N14}$ | 2581.6 | 2579.6 | -2 | |
| ACCCACCA-$K^{N15,N15}$ | 2583.6 | 2581.5 | -2 | |
| CCA-$K_9$ | 2033.7 | 1897.3 | -136.4 | $A_{base}$ (135.1)[1] |
| ACCA- $K_9$ | 2362.7 | 2273.4 | -89.3 | - $A_{base}$ (135.1)[1] –OH(17)[2] +$PO_2$ (63)[3, 4] |
| CCACCA- $K_9$ | 2972.7 | 2931.5 | -41.2 | $CO_2$ (44)[5]; |
| CCCACCA- $K_9$ | 3276.8 | 3189.5 | -87.3 | - $A_{base}$ (135.1)[1] –OH(17)[2] +$PO_2$ (63)[3, 4] |

Table S2. Theoretical and measured tandem MS/MS on 2273 (ACCA-$K_9$) and analysis. Only the $N^{14}$, $N^{14}$-lysine product 2273 MS/MS peaks are shown. The $N^{15}$,$N^{15}$-lysine products 2291 MS/MS peaks shift +18 relative those of un-labeled spectra.

| species | Theoretical Mass (M+1) | Measured Mass | Difference |
|---|---|---|---|
| 2273-$C_{base}$[1] | 2161.9 | 2162.2 | +0.3 |
| 2273-$A_{base}$[1] | 2137.9 | 2138.2 | +0.3 |
| 2273-A | 1943.8 | 1944.1 | +0.3 |
| 2273-AC | 1638.6 | 1638.2 | -0.4 |

Reference:

[1] Andersen, T. E., Kirpekar, F., and Haselmann, K. F. (2006) RNA fragmentation in MALDI mass spectrometry studied by H/D-exchange: mechanisms of general applicability to nucleic acids, *J Am Soc Mass Spectrom 17*, 1353-1368.

[2] Banerjee, S., and Mazumdar, S. (2012) Selective deletion of the internal lysine residue from the peptide sequence by collisional activation, *J Am Soc Mass Spectrom 23*, 1967-1980.

[3] Joyner, J. C., Keuper, K. D., and Cowan, J. A. (2013) Analysis of RNA cleavage by MALDI-TOF mass spectrometry, *Nucleic Acids Res 41*, e2.

[4] Ni, J., Pomerantz, C., Rozenski, J., Zhang, Y., and McCloskey, J. A. (1996) Interpretation of oligonucleotide mass spectra for determination of sequence using electrospray ionization and tandem mass spectrometry, *Anal Chem 68*, 1989-1999.

[5] Shek, P. Y., Zhao, J., Ke, Y., Siu, K. W., and Hopkinson, A. C. (2006) Fragmentations of protonated arginine, lysine and their methylated derivatives: concomitant losses of carbon monoxide or carbon dioxide and an amine, *J Phys Chem A 110*, 8282-8296.
